# Mobile nucleosomes and the accessibility of transcription factors to DNA: a modified dynamic equilibrium model

**DOI:** 10.1101/187815

**Authors:** Hadeel Khamis, Sergei Rudnizky, Ariel Kaplan

## Abstract

Nucleosomes, the basic building block of chromatin, regulate the accessibility of the transcription machinery to DNA. Recent studies have revealed that the nucleosome's spontaneous, thermally driven positional dynamics are modulated by different factors, and exploited by the cell as a regulatory mechanism. In particular, enrichment of mobile nucleosomes at the promoters of genes suggests that the mobility of nucleosomes may affect the ability of transcription factors to bind DNA. However, a quantitative model describing the effect nucleosome mobility on the effective affinity of transcription factors is lacking. We present here a simple equilibrium model that captures the essence of the effect, and show that modulation of the nucleosome's mobility can be a potent and versatile regulator of transcription factor binding.

## Introduction

DNA in our cells is packed into chromatin, a hierarchical structure of DNA and proteins whose basic building block is the nucleosome (Fig. 1A), composed of ~150 base-pairs (bp) of DNA wrapped around an octamer of histone proteins [1]. Packaging of the DNA, although essential, also presents a challenge for the cellular machinery in charge of reading the genetic information, since it reduces its accessibility to the DNA. As a result, modulating the local and global structure of chromatin is the most basic layer in the multi-layer regulation of gene expression [2].

**Figure 1:**
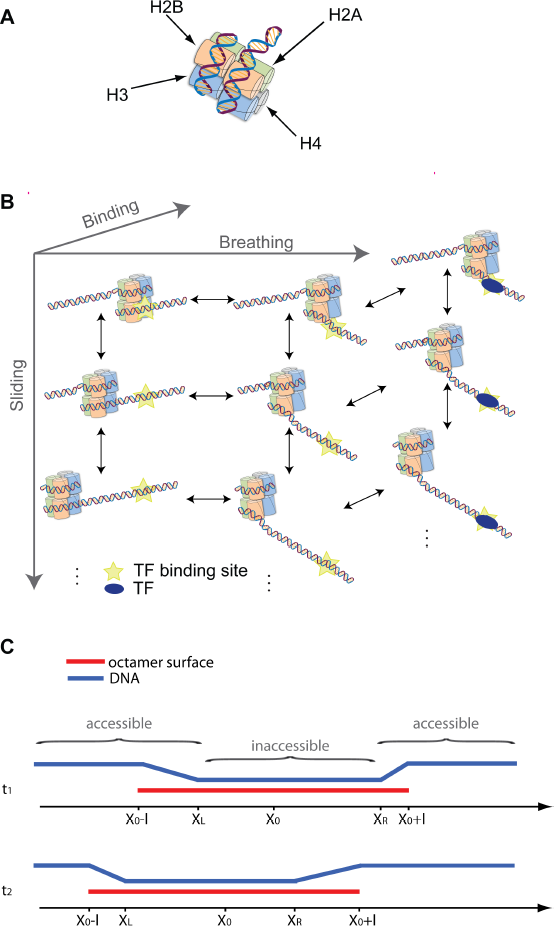
Nucleosome structure and dynamics. (A) Schematic structure of a nucleosome. ~147 bp of DNA wrapped around an octamer of histone proteins. (B) Transcription factors exploit a spontaneous breathing fluctuation (horizontal axis) that exposes their binding site, to bind to the DNA. In the presence of a mobile nucleosome, sliding (vertical axis) and breathing both modulate the accessibility to the DNA. (C) Schematic description of the variables used in the model.

Interestingly, while elucidating what determines the *mean* position of nucleosomes on genomic DNA has been the subject of numerous works [3–5], it is also clear that the *dynamics* of nucleosomes plays a crucial role in their function as regulators of DNA accessibility. Moreover, in addition to the intensely studied, long scale movements induced by ATP-consuming chromatin remodelers [6], understanding the role of thermally-driven conformational changes is crucial too. Two types of thermally driven conformational dynamics have been demonstrated: The first, comprising the spontaneous unwrapping of DNA at one end of the nucleosome (usually denoted nucleosome “breathing”), has been shown to play an important role in transcriptional initiation, more specifically on the ability of transcription factors (TFs) to bind DNA. By monitoring the ability of restriction enzymes to cut DNA at nucleosome-protected sites, it was shown [7] that TFs are able to DNA by exploiting these breathing fluctuations that momentarily expose their binding sites. Following these initial “bulk” studies, single-molecule FRET experiments directly detected the breathing fluctuations and studied their role in modulating TF accessibility [8–15]. Nucleosomal breathing has also been shown to affect the ability of RNA polymerase (RNAP) to transcribe through a nucleosome: Upon encountering the nucleosome, RNAP often backtracks allowing the reformation of disrupted DNA-octamer contacts [16]. Recovery from the backtracked state requires realignment of RNAP’s active with the 3’-end of the transcript [17]. Since, in the absence of polymerization, no source of chemical energy is available, the realignment can be achieved only by diffusion of RNAP on the DNA. However, the newly formed octamer-DNA contacts prevent the realignment. It has been shown [18] that recovery from the backtracked state can only take place by exploiting a spontaneous breathing fluctuation of the nucleosome, making breathing also an important factor in transcriptional elongation.

The second type of thermally driven dynamics present in the nucleosome is “thermal sliding” (also known as mobility), a spontaneous repositioning of the nucleosome where the histone octamer as a whole moves relative to the DNA. Early reports of sliding were based on nucleosomes reconstituted on 200-400 bp DNA fragments, and used as a reporter the electrophoretic mobility differences of complexes as a function of the octamer position of the DNA [19–21]. In other works, chemically modified histone proteins capable of inducing a nick in the DNA were used [22]. These studies revealed that nucleosomes reposition on their templates at time scales of hours, if incubated at 37°C, but not at 5°C. However, the relevance of these important findings to real genes was unclear since they were generally done with artificial, high affinity positioning sequences, such as Widom’s “601” sequence. In addition, they lacked the resolution to detect movements in the bp scale, and suffered from the limitations of bulk biochemical methods, with their intrinsic averaging over an unsynchronized population. Recently, we were able to probe the mobility of nucleosomes at the single molecule level, with bp-scale resolution, and on natural, biologically relevant sequences, using single molecule DNA unzipping with optical tweezers to probe the position of a single nucleosome several times as a function of time [23,24]. The single molecule experiments used as a model the promoters of *Cga* and *Lhb*, the genes that encode for the two subunits of the Luteinizing Hormone (LH), which are expressed under the same hormonal control but at levels that differ by as much as 7000-fold [23]. Surprisingly, we found that the mobility of nucleosomes located near the Transcription Start Site (TSS) in both genes, is higher than the mobility of nucleosomes downstream (+1 nucleosomes). Moreover, we found that histone variant H2A.Z, which, in cells expressing these genes, selectively replaces H2A at different positions (TSS nucleosome in Lhb and +1 nucleosome in Cga) also leads to a significantly higher mobility. Taken together, our previous results suggest that specifically modulating the mobility of nucleosomes is used by the cell as a regulatory mechanism.

Despite the experimental evidence suggesting the role of nucleosome thermal sliding in transcription, it is not yet clear how the mobility of nucleosomes affects the different phases of transcription. Here, we present a simple, equilibrium model for the effect of nucleosome mobility on the ability of TFs to bind to their binding site. Our model expands the dynamic equilibrium of Polach and Widom[7] by introducing, in addition of the thermal breathing, also the thermal sliding of nucleosomes.

## Results

The dynamic equilibrium model of Polach and Widom [7] postulates that binding of TFs to sites that are buried inside the nucleosome is modulated by the nucleosome’s thermally driven, spontaneous breathing. Breathing fluctuations are fast [9], and thus TFs bind to a buried site with an apparent dissociation constant

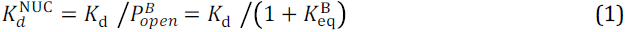

where 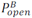 is the probability that the binding site is accessible following a breathing fluctuation, *K*_d_ is the dissociation constant of the TF on naked DNA, and 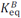 the equilibrium constant for a breathing fluctuation that exposes the binding site. The free-energy cost for such a fluctuation depends on the amount of DNA that needs to unwrap to expose the site. Hence, 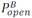 is a function of the distance of the binding site from the dyad, |*χ*_S_ − *χ*_0_|, where *χ*_0_ is the position of the dyad, and *χ_S_* is the position of the binding site. Note, that *χ_S_* can be on either side of the dyad, *χ*_0_ ≤ *χ_S_* or *χ*_0_ ≥ *χ_S_*. The total length of the nucleosome is defined as 2*l* = 147 bp. The conformational state of the nucleosome is characterized by three variables: its dyad position, *χ*_0_, and the position of the last wrapped bp on each side of the dyad, *χ_L_* and *χ_R_*, which can have any value in the intervals [*χ*_0_ − *l*, *χ*_0_] and [*χ*_0_, *χ*_0_ + *l*], respectively (Fig. 1C). Defining as *γ* < 0 the net interaction energy per bp, in units of *κ_B_T*, the free energy for a given *χ*_0_, and a given conformation can be written as 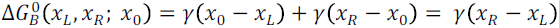. With this definition, the energy of a completely closed nucleosome is (*χ_R_* = *χ*_0_ + *l* and *χ_L_* = *χ*_0_ − *l*) is 2*γl* < 0, and the energy of a completely open on (*χ_R_* = *χ*_0_ and *χ_L_* = *χ*_0_) is 0.

The occupancy of a specific state characterized by *χ_L_* and *χ_R_* is given by 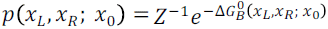, where 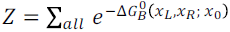 is the partition function and Σ_*all*_ indicates summation over all the possible states of the system. Replacing the sums with integrals, we can calculate *Z* as:

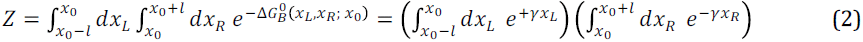

and therefore:

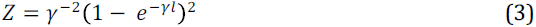

To find the probability, 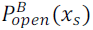, that the nucleosome is open to an extent that exposes *χ_S_*, we need to calculate *Σ_exposed_ p*(*χ_L_*, *χ_R_*; *χ*_0_, or 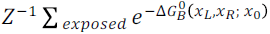, where *Σ_exposed_* is a summation of all the states for which the site at *χ_S_* is exposed, i.e. states for which *χ*_S_ > *χ_R_* or *χ*_S_ < *χ_L_*. Hence,

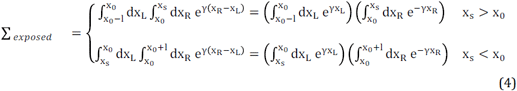

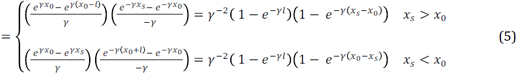

which can also be simply written as

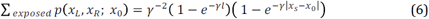

Combining Eqs. 3 and 6 results in:

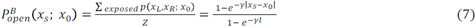

To generalize also for sites that are not covered by the nucleosome, i.e. |*χ*_S_ − *χ*_0_| > *l*, for which 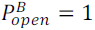, we write:

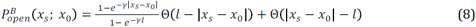

where Θ(x) is the step function. This recapitulates the classical results of Polach and Widom [7], and a later treatment by Prinsen and Schiessel [25]. Experiments with restriction enzymes confirmed this result, and demonstrated that *P_open_* is a sensitive function of the distance of the binding site from the nucleosome’s dyad, ranging from ~10^−2^-10^−1^ for sites at the edges of the nucleosome to ~10^−4^-10^−5^ for sites near to the dyad [7,9,26].

How should the mobility of a nucleosome be incorporated in such a model? Typical unwrapping and rewrapping rates are fast (~4 s^-1^ and ~20 to ~90 s^-1^, respectively [9]) as compared to the typical rates of repositioning; thus, we postulate a simple model in which breathing fluctuations are always in equilibrium for the instantaneous position *χ*_0_(*t*) of the nucleosome. In this case, we can assume that, on average, TF binding will be determined by a time-averaged accessibility, 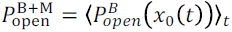, which now includes both breathing and mobility (Fig. 1B). To introduce the mobility in the model, we assume that at time 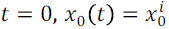, and model the nucleosome’s movement as a one-dimensional diffusion process, described by a probability distribution function given by: 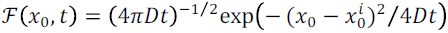, where D is the nucleosome’s diffusion constant. After a repositioning time *t*, the mean open probability 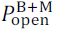 can be calculated as the expectation value of 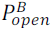, i.e. 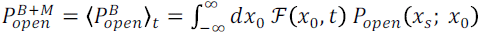 or

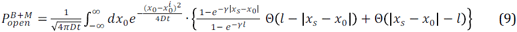

We show in the Appendix that, defining the dimensionless units 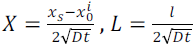 and 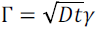, this results in the following expression:

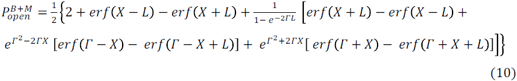

where *erf* is the error function. It is interesting to examine this expression at its limiting conditions. First, as expected, for *D* → 0 or *t* → 0 the equation reduces to the previous result of Eq. 8, when the nucleosome was static. Next, for a long time or high diffusion constant, 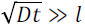, which implies *L* ≪ 1, Eq. 10 reduces to simply 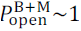, regardless of the position of the binding site. This reflects the fact that, on an infinite DNA molecule as we assumed in our model, averaging over a long enough time means that the “memory” of the initial localization is lost, and the nucleosome will spend equal amounts of time at all positions. So, the nucleosome will be most of the time far away from any specific single binding site and therefore all sites will be effectively accessible. This is clearly not a realistic scenario, since the motion will be limited in vivo by other factors, e.g. neighboring nucleosomes. However, it highlights the physical effect of nucleosome mobility on the accessibility: motion of the nucleosome will tend to reduce, with time, the differences in accessibility by different sites. This can be intuitively understood by considering that, if a binding site is outside the nucleosome, but in its vicinity, 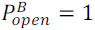. Hence, the mobility can only have a repressing effect, as the mobile nucleosome will be able to cover the binding site, resulting in 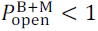. Alternatively, if the binding site close to the dyad, 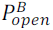 has its minimal possible value, hence repositioning can only increase the exposure, i.e. 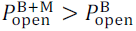. This is also shown in Fig. 2A, where the results from Eq. 10 are plotted for different times, and for values that were derived from measurements on real nucleosomes: γ=-0.1 k_B_T/bp (Ref. [25]) and D=1.5 bp^2^/s (Ref. [24]). Interestingly, the fact that the accessibility of certain sites is increased, while the accessibility of others is decreased, means that the modulation by the mobility can have both a repressing as well as a facilitating effect on transcription. Fig. 2B shows how that modulating the nucleosomes diffusion constant can have a significant effect: increasing D by a factor of 2, a change of similar magnitude to the differences in mobility observed by the introduction of H2A.Z [23], results, over a time of 3 min, in a site-specific increase in the accessibility that can be as high as 4-fold.

**Figure 2:**
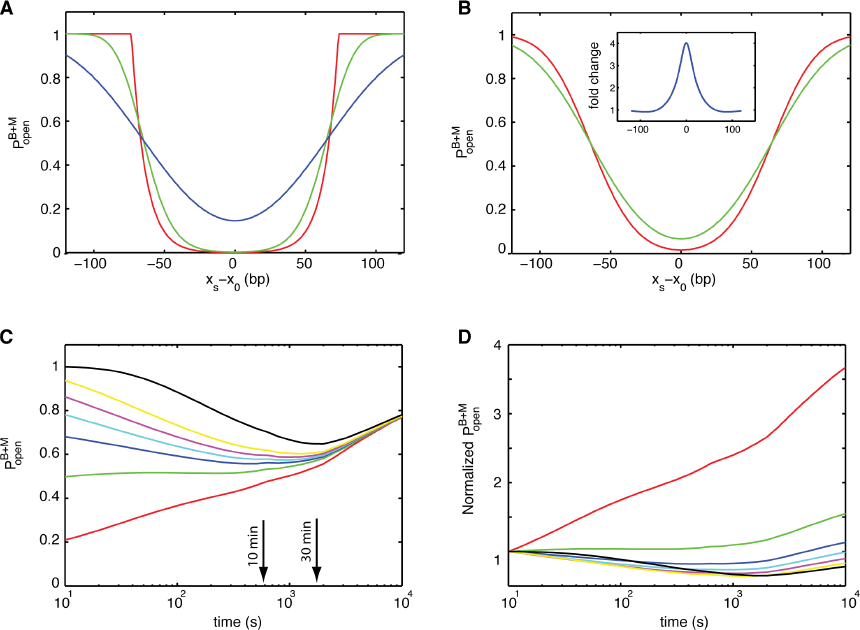
Nucleosomal mobility modulates the accessibility of TF to DNA. (A) The probability for a binding site to be exposed, as a function of its distance from the nucleosome’s dyad mean position. Shown are the probabilities for t=0 (red), 60 (green) and 600 (blue) s. The curves correspond to D=1.5 bp^2^/s and γ=-0.1 k_B_T/bp. (B) Probabilities for two nucleosomes with different mobilities, D=1.5 (red) and 3 (green) bp^2^/s. Shown are results for t=180 s and γ=-0.1 k_B_T/bp. Inset: ratio of the probabilities. (C) Temporal dynamics of site exposure. Exposure probability as a function of time, for sites located at 55 (red), 65 (green), 70 (blue), 73 (cyan), 76 (magenta), 80 (yellow) and 95 (black) bp from the dyad’s mean position. The curves correspond to D=3 bp^2^/s and γ=-0.1 k_B_T/bp. (D) Fold-change in the exposure probability, relative to t=0. Same data as in (A).

In general, sites that are closer to the dyad than a critical value will have their accessibility increased, while those that are further away than this value will see a decrease in accessibility. However, the position of this critical value shifts away from the dyad’s mean position with time, so positions that saw initially a decrease in accessibility will later on experience an increase in it. Hence, our model also predicts that the mobility can be used to produce a time-dependent accessibility. Figs. 2C,D show the accessibility of 7 different sites, ranging from 55 to 95 bp from the dyad’s mean position. As shown in the figure, depending on the specific position, a rich variety of temporal profiles can be observed, including a monotonic increase, and a temporary repression of different levels.

## Discussion

A mechanistic understanding of transcriptional regulation requires characterizing how different factors, such as the sequence of DNA, the identity of the histone proteins, post- translational modifications, chromatin remodelers and distal enhancers, affect the biophysical properties of nucleosomes (e.g. their position, stability and mobility), and how these properties in turn affect the transcriptional outcome.

We presented here a simple, equilibrium model for the ability of TF to bind their target site. Building on previous results, we incorporated the thermal mobility of nucleosomes by modeling this motion as a simple 1D diffusion process. The results from this model are intuitive: the movement of the nucleosome tends to moderate the large differences in accessibility between sites buried deep inside the nucleosome and those that are initially not affected by its presence. This is analogous to the tendency of any diffusional process towards a uniform state. However, the implications of this model are far reaching for the regulation of genes: Our results highlight how the mobility of nucleosomes can affect the differential accessibility of different sites, and the differential accessibility of the same site before and after the incorporation of a nucleosome with a different mobility. Moreover, they show how different patterns of time-dependent accessibility can be achieved for different sites and different nucleosomes. Taken together, these results make the modulation of the mobility a powerful and versatile tool that provides a way to moderately adjust TF binding, as opposed to the more radical effect of eviction of the nucleosome.

## Acknowledgements

We acknowledge support from the Israel Science Foundation (Grant 1782/17), the Israeli Centers of Research Excellence program (I-CORE, Center no. 1902/12), the European Commission (Grant 293923), the Eliyahu Pen Research Fund and the J. S. Frankford Research Fund.

## Appendix 1

In order to calculate

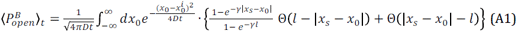

we split the integral’s integration range as, i.e. 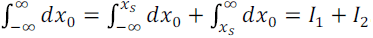. Hence,

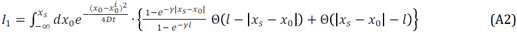

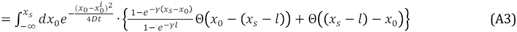

The integral can be split again as

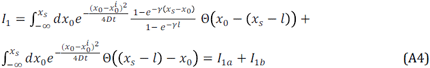

We now eliminate the step function by modifying the limits of the integrals:

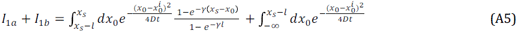

In the same way, for *I*_2_,

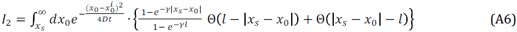

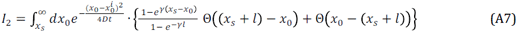

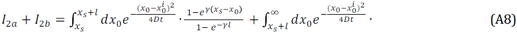

Hence, the integrals we need to solve are:

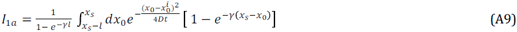

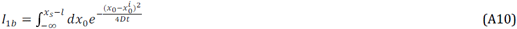

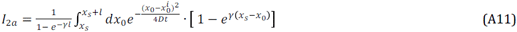

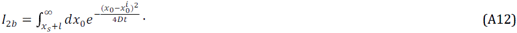

and their solutions are

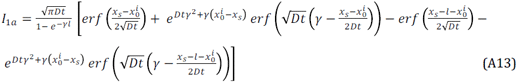

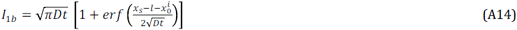

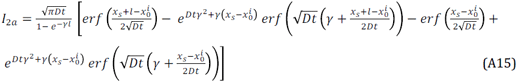

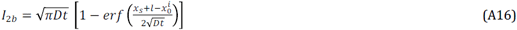

Next, we define dimensionless units: 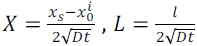 and 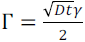. With them, the sum of the four integrals is:

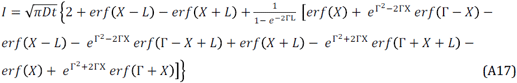

and therefore:

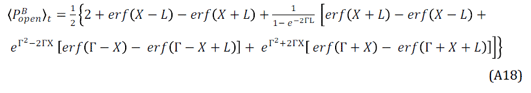

## References

[1] K. Luger, a W. Mäder, R. K. Richmond, D. F. Sargent, and T. J. Richmond, Nature (1997).

[2] O. Bell, V. K. Tiwari, N. H. Thomä, and D. Schübeler, Nat. Rev. Genet. 12, 554 (2011).

[3] N. Kaplan, I. K. Moore, Y. Fondufe-Mittendorf, A. J. Gossett, D. Tillo, Y. Field, E. M. LeProust, T. R. Hughes, J. D. Lieb, J. Widom, and E. Segal, Nature 458, 362 (2009).

[4] Y. Zhang, Z. Moqtaderi, B. P. Rattner, G. Euskirchen, M. Snyder, J. T. Kadonaga, X. S. Liu, and K. Struhl, Nat. Struct. Mol. Biol. 17, 920 (2010).

[5] K. Struhl and E. Segal, 20, 267 (2013).

[6] C. R. Clapier, J. Iwasa, B. R. Cairns, and C. L. Peterson, Nat. Rev. Mol. Cell Biol. 18, 407 (2017).

[7] K. J. Polach and J. Widom, J. Mol. Biol. 254, 130 (1995).

[8] G. Li and J. Widom, Nat. Struct. Mol. Biol. 11, 763 (2004).

[9] G. Li, M. Levitus, C. Bustamante, and J. Widom, Nat. Struct. Mol. Biol. 12, 46 (2005).

[10] M. Tomschik, H. Zheng, K. Van Holde, J. Zlatanova, and S. H. Leuba, (n.d.).

[11] ‡ L. Kelbauskas, ‡,§ N. Chan, ‡,§,‖ R. Bash, ⊥ J. Yodh, ‡,§ and N. Woodbury, and § D. Lohr*, (2007).

[12] L. Kelbauskas, J. Sun, N. Woodbury, and D. Lohr, Biochemistry 47, 9627 (2008).

[13] A. Gansen, K. Tóth, N. Schwarz, and J. Langowski, J. Phys. Chem. B 113, 2604 (2009).

[14] A. Gansen, A. Valeri, F. Hauger, S. Felekyan, S. Kalinin, K. To, C. A. M. Seidel, K. Toth, J. Langowski, and C. A. M. Seidel, Proc. Natl. Acad. Sci. U. S. A. 106, 15308 (2009).

[15] W. J. A. Koopmans, R. Buning, T. Schmidt, and J. Van Noort, (n.d.).

[16] D. A. Gaykalova, O. I. Kulaeva, O. Volokh, A. K. Shaytan, F.-K. Hsieh, M. P. Kirpichnikov, O. S. Sokolova, and V. M. Studitsky, Proc. Natl. Acad. Sci. U. S. A. 112, E5787 (2015).

[17] E. A. Galburt, S. W. Grill, A. Wiedmann, L. Lubkowska, J. Choy, E. Nogales, M. Kashlev, and C. Bustamante, Nature 446, 820 (2007).

[18] C. Hodges, L. Bintu, L. Lubkowska, M. Kashlev, and C. Bustamante, Science 325, 626 (2009).

[19] S. Pennings, G. Meersseman, and E. M. Bradbury, J. Mol. Biol. 220, 101 (1991).

[20] G. Meersseman, S. Pennings’, and E. M. Bradbury’, EMBO J. 1, 2951 (1992).

[21] S. Pennings, G. Meersseman, and E. M. Bradbury, Proc. Natl. Acad. Sci. U. S. A. 91, 10275 (1994).

[22] A. Flaus and T. J. Richmond, J. Mol. Biol. 275, 427 (1998).

[23] S. Rudnizky, A. Bavly, O. Malik, L. Pnueli, P. Melamed, and A. Kaplan, Nat. Commun. 7, 12958 (2016).

[24] S. Rudnizky, O. Malik, A. Bavly, L. Pnueli, P. Melamed, and A. Kaplan, Protein Sci. 26, 1266 (2017).

[25] P. Prinsen and H. Schiessel, Biochimie 92, 1722 (2010).

[26] J. Anderson and J. Widom, J. Mol. Biol. 296, 979 (2000).

